# Contact toxicity, antifeedant activity and oviposition preference of osthole against agricultural pests

**DOI:** 10.1101/2023.06.15.545046

**Authors:** Fang Dong, Xin Chen, Men Xingyuan, Zhuo Li, Yujun Kong, Yiyang Yuan, Feng Ge

## Abstract

Osthole, the dominant bioactive constituent in *Cnidium monnieri*, has been shown to exhibit acute insecticidal activities. However, its detailed toxicity, antifeedant and oviposition preference effects against agricultural pests has not been fully understood, which has greatly hindered its practical applications. This study is designed to investigate the contact toxicity, antifeedant activity and oviposition preference of osthole against three agricultural pests (*T. urticae*, *M. persicae* and *B. dorsalis*) to evaluate its potential agricultural applications. Our results showed that *Cnidium monnieri* (L.) Cusson (CMC) have a high osthole content of 11.4 mg/g. Osthole exhibited a comparable level of acute toxicity against *T. urticae* to four other coumarins found in CMC. Osthole demonstrated significant insecticidal activity against first instar nymphs and adults of *T. urticae* and *M. persicae* in a dose-dependent manner, but not against *B. dorsalis* adults. Osthole exposure reduced the fecundity and prolonged the developmental time of *T. urticae* and *M. persicae*. Leaf choice bioassays revealed potent antifeedant activity in *T. urticae*. Furthermore, female *B. dorsalis* showed a distinct preference for laying eggs in mango juice with 0.02 mg/mL osthole at 48 hours, a preference which persisted at 96 hours. These results provide valuable insights into the toxicity, repellent activity, and attractant activity of osthole, thereby contributing to its expanded use in pest control.

## 1 Introduction

The intensive use of chemical insecticides has led to the development of pesticide resistance, ecosystem imbalances, environmental deterioration and health risks (Carvhalo, 2017; Tudi et al., 2021). The botanical insecticides, with reduced health and environmental impacts, are promising alternatives to chemical insecticides (Isman, 2006; Isman, 2020).

*Cnidium monnieri* is a medicinal plant that is mainly grown in China, Japan, Korea and Vietnam (https://www.gbif.org/species/3034720). *Cnidium monnieri* (L.) Cusson (CMC), the dry fruit of *C. monnieri*, is commonly used as traditional Chinese medicine with a broad range of pharmaceutical properties (Li et al., 2015). CMC contains 429 chemical constituents including coumarins, volatile constituents, liposoluble compounds, chromones, monoterpenoid glucosides, terpenoids, glycosides and glucides (Sun et al., 2020). Coumarins are major chemical compounds in CMC, among which osthole is the dominant bioactive constituent with antitumor, anti-inflammatory, neuroprotective, osteogenic, cardiovascular protective, antimicrobial and antiparasitic activities (Cai et al., 2000; Liu et al., 1999; Sun et al., 2021; Wu et al., 2017).

Osthole (7-methoxy-8-(3-methyl-2-butenyl) coumarin), a simple coumarin, is mainly found in the genera of Umbelliferae and Rutaceae (Yan et al., 2012; Zhang et al., 2002). Recent research has highlighted its potential as an acute insecticide, displaying efficacy against green peach aphids (*Myzus persicae*) and two-spotted spider mites (*Tetranychus urticae*) (Yan et al., 2021). However, its detailed toxicity, antifeedant and oviposition preference effects against agricultural pests remain incompletely understood, significantly limiting its practical application.

*T. urticae*, *M. persicae* and oriental fruit fly (*Bactrocera dorsalis*) are cosmopolitan agricultural pests that feeds on hundred kinds of fruit and vegetables (Attia et al., 2013; Margaritopoulos et al., 2000; Paini et al., 2016). They can destroy many economically valuable host plants in agriculture and horticulture, including tomatoes, peppers, cucumbers, strawberries, maize, soy, apples, bananas, mango, grapes and citrus (Clarke et al., 2005; Helle and Sabelis, 1985; van Emden and Harrington, 2017). Because chemical insecticides are intensively used for controlling these pest insects, a broad cross-resistance within and between pesticide classes has been rapidly developed (Bass et al., 2014; Jin et al., 2011; Khajehali et al., 2011; Singh et al., 2021; Van Leeuwen et al., 2010; Wei et al., 2019). Therefore, the effective botanical pesticides are required as alternatives to conventional pesticides in the control of these pests. In this study, the efficacy of osthole was assessed for its contact toxicity, antifeedant activity and oviposition preference against *T. urticae*, *M. persicae* and *B. dorsalis* to determine its potential agricultural applications.

## 2 Materials and methods

### 2.1 Insects and reagents

A colony of *T. urticae* was collected from a strawberry field population during November 2021 in Shandong Province, China, and subsequently raised on caged common-bean (*Phaseolus vulgaris*). The *M. persicae* (green clone) used in this study were derived from a wild population collected in the oilseed rape (*Brassica napus*) fields of Shandong province, China, during March 2022. Aphids were reared under laboratory conditions on caged Chinese cabbage (*B. campestris*). *B. dorsalis*, which were originally obtained from a continuously maintained culture at the Institute of Plant Protection, Shandong Academy of Agricultural Sciences, were maintained by hatching larvae on bananas and rearing adult flies in cages with an artificial diet consisting of yeast extract and sugar. All insects were maintained under controlled conditions at 25 ± 1 °C with a photoperiod of 16:8 hours (L:D) until use in the experiments.

CMC was purchased from Binnong Technology Co., Ltd (Wudi, Shandong), while osthole with a purity of 99.5%, and Dimethylsulfoxide (DMSO) and Triton X-100 were respectively obtained from the National Institutes for Food and Drug Control, Sinopharm Chemical Reagent Co., Ltd, and Solarbio Co., Ltd. Reference compounds used in this study included imperatorin (98%), methoxsalen (99.98%), xanthotoxol (98%), columbianadin (98%), isopimpinellin (98%), 5-methoxypsoralen (98%), meranzin (hydrate) (98%), angelicin (98%), auraptenol (98%), isogosferol (97%) and scoparone (98%), which were purchased from MedChemExpress (Shanghai), Aladdin (Shanghai), Yuanye (Shanghai), Acmec (Shanghai), Glpbio (Montclair), Bidepharm (Shanghai), Macklin (Shanghai) and Desite (Chengdu), respectively.

### 2.2 Extraction and quantification of coumarins in CMC

Coumarins were extracted from CMC using a modified method based on Bourgaud et al., 1994. In brief, 10 g of powdered CMC was placed in sterile medical gauze and extracted three times with 100 mL of 95% ethanol using a Soxhlet extractor for 3.5 h each time. The resulting extracts were combined, filtered, and concentrated under vacuum at 50 °C. The concentrate was then dissolved in 25 mL of methanol and subjected to centrifugation at 10,000 rpm at 4 °C for 5 minutes. A 250 μL aliquot of the supernatant was transferred to a new tube and made up to 25 mL with methanol. Before chromatographic analysis, the solution was filtered through a 0.22 μm PTFE membrane.

The quantification of auraptenol, columbianadin, xanthotoxol, and isopimpinellin was performed using a Waters ACQUITY UPLC I-Class liquid chromatography (LC) system coupled to a TQ-S triple quadrupole mass spectrometer (MS) with an electrospray ionization (ESI) source (Waters Corp., Milford, MA, USA). Chromatographic separation was carried out on a Waters AcQuity UPLC BEH C18 column (4.6 mm × 150 mm, 1.7 μm) with an injection volume of 2 μL for all samples and a mobile phase flow rate of 5 μL/min. The gradient elution was performed using 0.1% fomic acid (A) and acetonitrile (B) with the following conditions: 0-0.5 min, 30% B; 0.5-1 min, 30% B to 90% B; 1-4 min, 90% B; 4-5 min, 90% B - 30% B. Quantification was conducted in positive ion multiple reaction monitoring (MRM) mode with a capillary voltage of 3.0 kV, desolvation temperature of 500 °C, and flow rates of 850 and 150 L/h for desolvation and cone gas, respectively.

Osthole, imperatorin, isopimpinellin, 5-methoxypsoralen, methoxsalen, meranzin (hydrate), angelicin, and isogosferol were quantified using a Shimadzu Nexera X2 LC-30AD liquid chromatography (LC) system coupled to an AB SCIEX TRIPLE QUAD 4500 mass spectrometer (MS) with an electrospray ionization (ESI) source (Shimadzu Corp., Chiyoda-ku, Tokyo, Japan). Chromatographic separation was carried out on a Angilent Poroshell 120 EC C18 column (4.6 mm × 100 mm, 2.7 μm) with an injection volume of 5 μL and a mobile phase flow rate of 0.4 mL/min. A gradient elution consisting of 0.1% formic acid (A) and Methanol (B) was used to separate the coumarins, with the following gradient conditions: 0-0.2 min, 10% B; 0.2-2 min, 10% B to 90% B; 2-8 min, 90% B; 8-8.1 min, 90% B - 10% B. Quantification was performed in positive ion multiple reaction monitoring (MRM) mode with two MRM transitions, Q1 (precursor ion) and Q3 (product ion), and a dwell time of 50 ms for quantitation and identification of the coumarins based on mass-to-charge ratio (m/z) calculations.

### 2.3 Acute toxicity bioassay

The acute toxicity bioassays of osthole and other coumarins against *T. urticae* and *M. persicae* were conducted using a previously described leaf-dipping method with some modifications (Mostafiz et al., 2020). Osthole was first dissolved in DMSO and then diluted with distilled water containing 0.1% Triton X-100 to varying concentrations, with the final DMSO concentration being 3% in the final dilutions. Individual 60 mm cut leaf discs, each with 20 one-day-old adult females, were dipped in different osthole dilutions for a duration of either 5 s (for *T. urticae*) or 10 s (for *M. persicae*). The excess solution was then blotted off with filter papers, and the insects were subsequently transferred to an agar-containing Petri dish with a new leaf disc. Insects treated with distilled water containing 0.1% Triton X-100 and 3% DMSO via the same method served as the control. Mortality was recorded every 24 h, and each treatment was repeated five times. The larvicidal bioassays were conducted using a similar methodology, except leaf discs infested with 1^st^ instar nymphs were immersed in osthole solutions for a duration of 10 seconds. Mortality was assessed 3 days’ post-experiment, and each treatment was repeated five times. The insecticidal activities of osthole against *B. dorsalis* were evaluated using a modified feeding method (Zhang et al., 2015). Batches of 20 three-day-old adult flies, consisting of 10 females and 10 males, were subjected to a 6-hour starvation period followed by feeding with artificial diets containing osthole. Mortality was assessed every 24 h over 7 days. Each treatment was repeated three times.

The LC_20_, LC_50_, and LC_90_ values were calculated using Probit analysis in SPSS 14.0 software. The LC_20_ and LC_50_ concentrations were subsequently used in experiments to evaluate the effects of osthole on larval growth and adult reproduction.

### 2.4 Larval growth and development bioassays

To evaluate the impact of osthole on larval growth and development, newly hatched mite or aphid larvae were immersed in the osthole dilutions (LC_20_ and LC_50_) using the aforementioned method. Distilled water containing 0.1% Triton X-100 and 3% DMSO was used as the control. Once 40 larvae that survived the treatment were obtained, they were individually reared in agar-containing Petri dishes with a fresh leaf disc. The duration of different developmental stages was recorded, and body lengths were measured one day after eclosion.

### 2.5 Fecundity bioassays

Newly emerged gravid female adults of *T. urticae* and *M. persica* were subjected to the osthole dilutions (LC_20_ and LC_50_) using the aforementioned leaf-dipping method. After 24 hours, the surviving females were individually transferred to agar-containing Petri dishes with a fresh leaf disc. The eggs of *T. urticae* and the nymphs of *M. persicae* were counted and removed daily for 3 days, with ten replicates carried out for each assay.

Meanwhile, the fecundity assay for *B. dorsalis* was conducted using a feeding method with some modifications (Li et al., 2022). Specifically, ten-day-old female and male flies (10 each) were placed in a plastic cup (250 mL) and fed with artificial feed containing osthole at different concentrations (0.0625-2 mg/mL). The control treatment involved artificial feed without osthole. The eggs were collected with a plastic tube (10 mL) along with mango juice, and the number of eggs was counted daily for 15 days. Each treatment was repeated ten times.

### 2.6 Antifeedant bioassays

To evaluate the antifeedant activities of osthole on *T. urticae*, we applied osthole to one half of a 35mm diameter common bean leaf disc using a sterile cotton swab, while the other half was treated with distilled water containing 0.1% triton X-100 and 3% DMSO as a control. After the solution dried, the leaf was transferred to a Petri dish and 30 female adult mites were introduced onto the central line, allowing aphids to choose between treated and untreated control areas. The number of mites on each half of the leaf area was recorded at 0.5, 1, 2, 6, 12, 24, and 48 hour intervals. Each concentration was tested with 30 mites, and eight replicates were performed.

To assess the antifeedant properties of osthole against *M. persicae*, Chinese cabbage leaves were briefly immersed in an osthole solution for 10 seconds, while similarly sized leaves were submerged in distilled water containing 0.1% triton X-100 and 3% DMSO as a control. The treated and control leaves were transferred to plastic boxes (10 × 16 cm) containing a 1 cm thick hydrophilic sponge, a 1 mm thick filter paper, and black PVC. Cohorts of one-day-old adult female aphids were introduced onto the central axis that bisected the experimental arena into two equal halves. The number of aphids on each leaf was recorded at 0.5, 1, 2, 6, 12, 24, and 48 hour intervals. Each concentration was tested on 30 aphids with eight replicates.

### 2.7 Oviposition preference bioassays

The oviposition preference of *B. dorsalis* was assessed using a modified version of the methodology described by Li et al. (2020). Egg collection devices were created by perforating 180 mL plastic cups with ten rows of ten holes each, positioned 5 cm from the bottom of the cup. The diameter of each hole was approximately 1 mm, with a spacing of about 0.5 cm between them. Each row was spaced at around 1.3 cm intervals. Two devices were placed in a sterilized cage (60 × 35 × 35 cm) containing 15 female and 15 male flies to inoculate eggs. One device contained a solution of osthole in mango juice with 3% DMSO, while the other served as a control and contained only mango juice solution with 3% DMSO. The surface of each device was also sprayed with the corresponding mango juice solutions. The number of eggs was recorded at 24, 48 and 72 hour intervals. The experiment was repeated eight times for each treatment.

### 2.8 Data analysis

We used ANOVA and Turkey’s HSD multiple comparison tests to determine differences in coumarin content in CMC and mortality rates of nymphs of *T. urticae* and *M. persicae* after exposure to osthole at different concentrations. ANOVA and Turkey’s HSD tests were also used to identify differences in the osthole’s impact on developmental period, body length, and reproduction. The paired sample T-test was employed to analyze antifeedant activity of osthole on female adults of *T. urticae* and *M. persicae*, as well as differences in oviposition attraction of osthole on female adults of *B. dorsalis*. All analyses were performed in GraphPad Prism 9.

## 3 Results

### 3.1 Osthole is the primary coumarin in CMC

Based on LC/MS analysis of CMC, osthole was found to be the most abundant coumarin in CMC, accounting for 67.1% of the total 11 coumarins determined, followed by imperatorin and xanthotoxol (Fig. 1). The contents of isopimpinellin, 5-methoxypsoralen, methoxsalen, meranzin (hydrate), auraptenol, and columbianadin ranged from 0.12 to 3.41 mg/g (Fig. 1). CMC only contains trace amounts of angelicin and isogosferol (Fig. 1).

**Figure 1.**
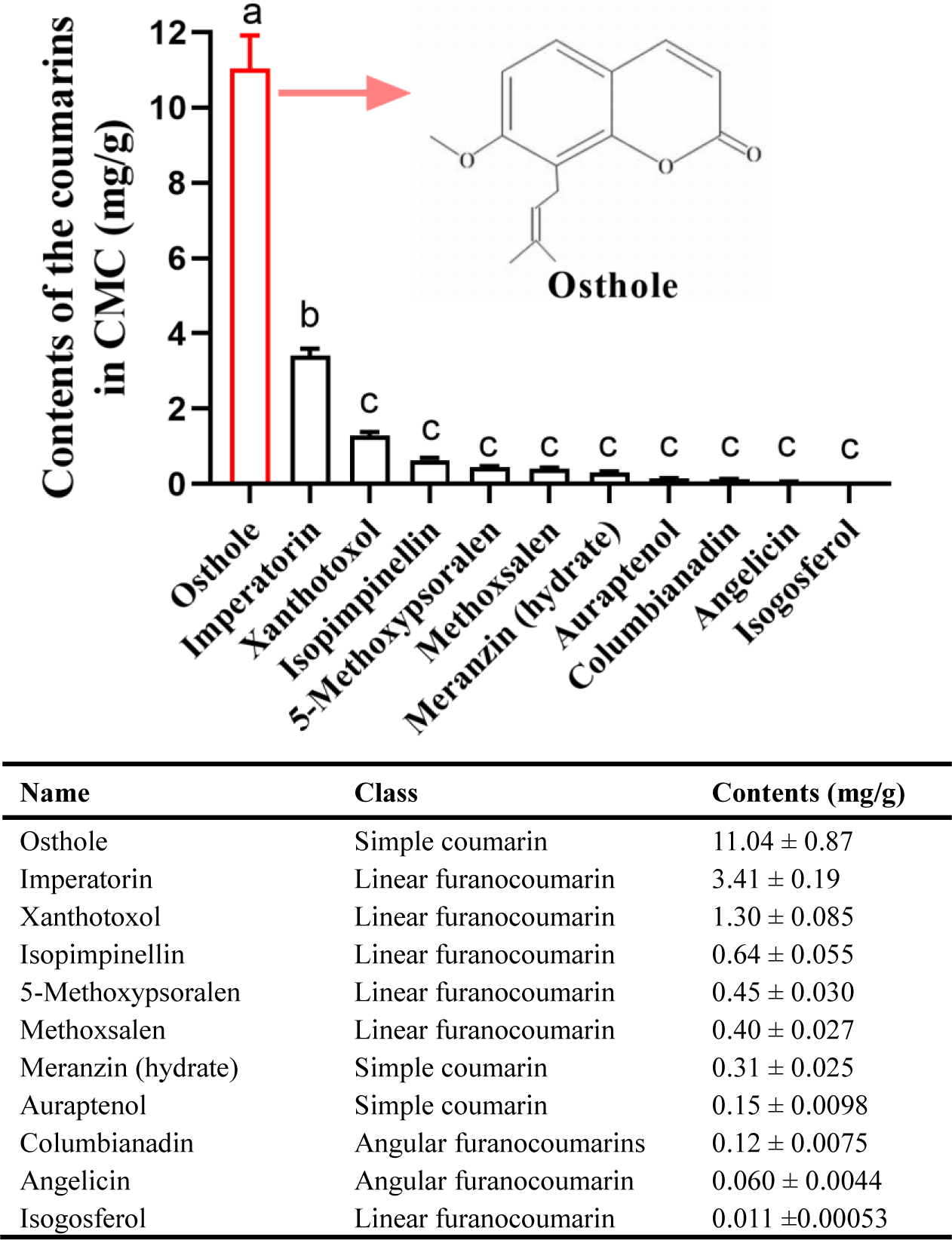
Contents of the coumarins in CMC. Different lowercase letters indicate significant differences among coumarins (one-way analysis of variance (ANOVA), followed by multiple comparisons of Tukey’s test, P < 0.05). Error bars represent standard error (n = 4).

### 3.2 Osthole showed comparable contact toxicity to other coumarins in CMC against *T. urticae*

We investigated the contact toxicity of 6 coumarins in CMC against adult *T. urticae* using a consistent concentration of 2 mg/mL and the leaf-dipping method, with scoparone as the positive control. Results revealed mortality rates of 23% to 42% across all 6 coumarins. Osthole caused higher mortality than the other coumarins, although statistical significance was not observed (Table 2).

**Table 1.** Efficacy of osthole against female adults of *T. urticae*.

| Time (d) | Slope ±<br>SE | Chi-square | *p*-Value | LC<sub>20</sub> (95% FL)<sup>a</sup> | LC<sub>50</sub> (95% FL)<sup>b</sup> |
| --- | --- | --- | --- | --- | --- |
| 1 | 1.44 ± 0.18 | 14.56 | 0.002 | 1.14 (0.001~2.56) | 4.39 (1.65~62.77) |
| 2 | 1.33 ± 0.19 | 9.52 | 0.023 | 0.87 (0.006~1.99) | 3.76 (1.43~17.00) |
| 3 | 1.41 ± 0.19 | 7.27 | 0.064 | 0.86 (0.054~1.77) | 3.39 (1.58~8.12) |
| 4 | 1.50 ± 0.19 | 4.24 | 0.237 | 0.79 (0.46~1.12) | 2.90 (2.27~3.67) |
| 5 | 1.41 ± 0.19 | 6.03 | 0.110 | 0.63 (0.047~1.33) | 2.49 (1.08~4.72) |

<sup>a</sup> Concentration of osthole killing 20% and its 95% fiducial limits in mg toxin per mL solution, n = 20 nymphs for each LC<sub>20</sub>.
<sup>b</sup> Concentration of osthole killing 50% and its 95% fiducial limits in mg toxin per mL solution, n = 20 nymphs for each LC<sub>50</sub>.

**Table 2.** Efficacy of osthole against female adults of *M. persicae*.

| Time (d) | Slope ± SE | Chi-square | *p*-Value | LC20 (95% FL)<sup>a</sup> | LC50 (95% FL)<sup>b</sup> |
| --- | --- | --- | --- | --- | --- |
| 1 | 2.07 ± 0.21 | 7.85 | 0.049 | 1.79 (0.69~1.99) | 4.56 (2.99~8.01) |
| 2 | 2.08 ± 0.20 | 15.67 | 0.001 | 1.46 (0.17~2.64) | 3.70 (1.80~8.70) |
| 3 | 2.33 ± 0.22 | 25.14 | 0.000 | 1.37 (0.002~2.77) | 3.15 (0.58~10.20) |
| 4 | 2.36 ± 0.22 | 23.19 | 0.000 | 1.31 (0.008~2.60) | 2.98 (0.67~7.93) |
| 5 | 2.29 ± 0.22 | 19.50 | 0.000 | 1.27 (0.034~2.45) | 2.95 (0.90~6.82) |

<sup>a</sup> Concentration of osthole killing 20% and its 95% fiducial limits in mg toxin per mL solution, n = 20 nymphs for each LC<sub>20</sub>.
<sup>b</sup> Concentration of osthole killing 50% and its 95% fiducial limits in mg toxin per mL solution, n = 20 nymphs for each LC<sub>50</sub>.

### 3.3 Osthole exhibits toxicity towards *T. urticae* and *M. persicae*, but not *B. dorsalis*

We tested osthole’s insecticidal activity against *T. urticae* and *M. persicae* and generated Kaplan-Meier 5-day survival curves to depict the daily survival rates of female adults exposed to osthole concentrations ranging from 0 to 10 mg/mL (Fig. 3A). Results showed a significant reduction in the survival rates of *T. urticae* and *M. persicae* with concentrations above 1.25 and 2.5 mg/mL, respectively, with the lowest survival rates observed on day 5 at 10 mg/mL for *T. urticae* and 5 mg/mL for *M. persicae* (Fig. 3A). The LC_20_ and LC_50_ values of *T. urticae* and *M. persicae* were both dependent on the time after immersion, indicating the delayed mortality effect of osthole, as shown in Table 1 and 2.

**Figure 2.**
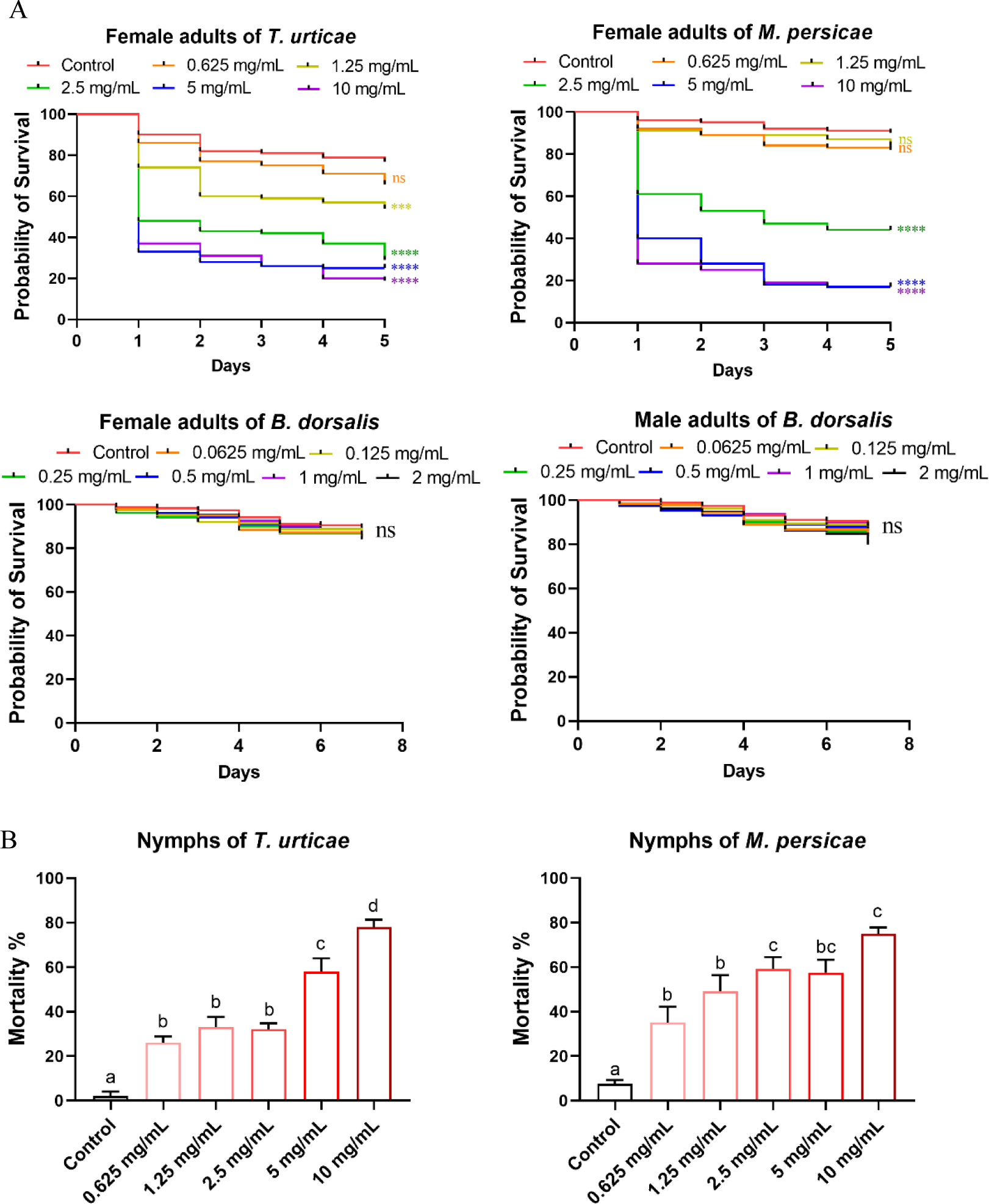
The acute toxicity of osthole against *T. urticae*, *M. persicae* and *B. dorsalis*. (A) Kaplan-Meier survival curve of adults from the three pest species upon exposure to osthole. For each treatment, 20 individuals were randomly assigned to 5 replicates. The asterisk indicates a statistically significant difference measured by using Log-Rank test as compared to the wild type (***, P < 0.0005; ****, P < 0.0001). (B) Mortality of *T. urticae* and *M. persicae* after osthole exposure (n = 20). Different lowercase letters indicate significant differences among coumarins (one-way analysis of variance (ANOVA), followed by multiple comparisons of Tukey’s test, P < 0.05). Error bars represent standard error.

**Figure 3.**
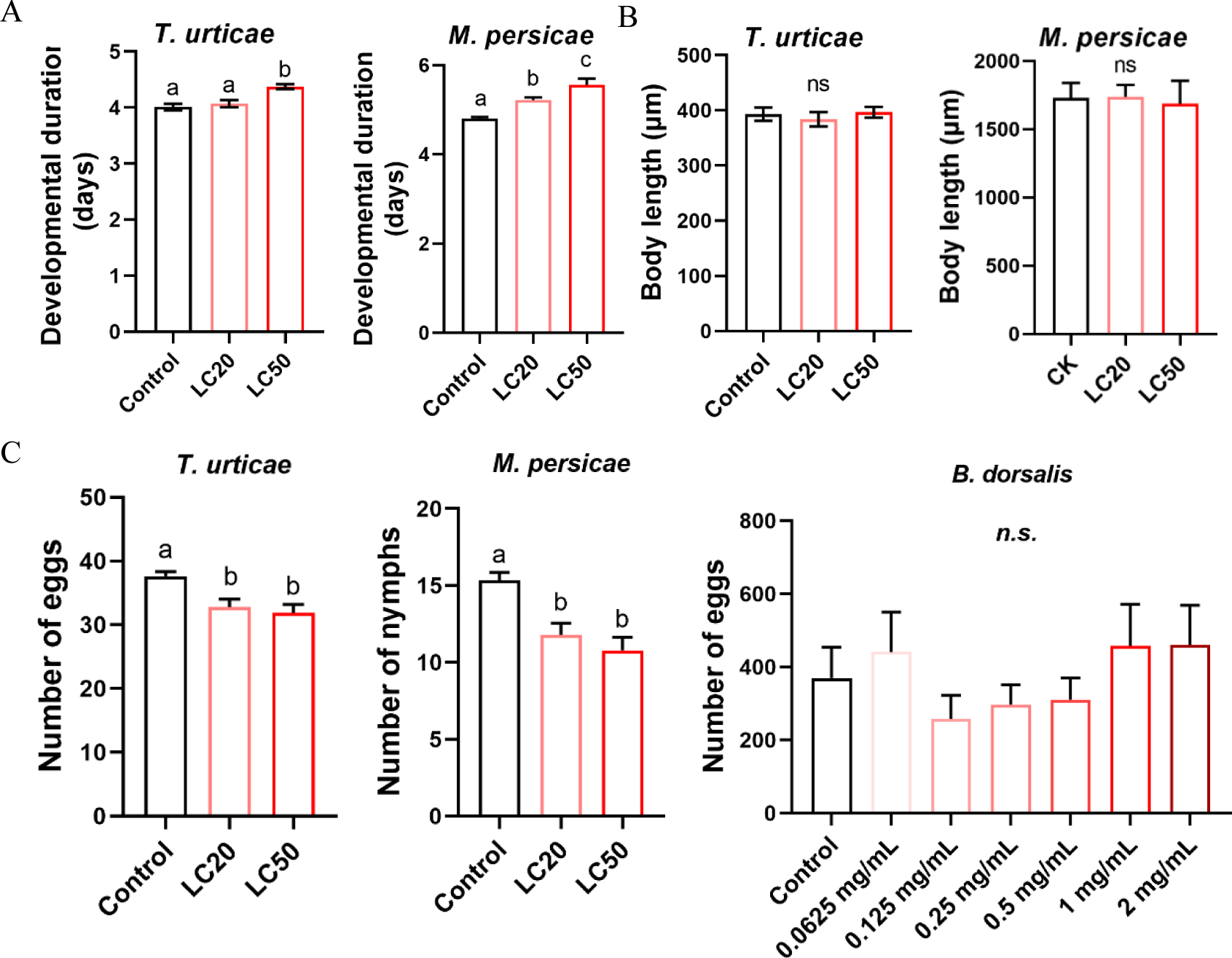
Effects of osthole treatments on the developmental period (A), body length (B) and reproduction (C) of *T. urticae* and *M. persicae*. Different lowercase letters indicate significant differences among treatments (one-way analysis of variance (ANOVA), followed by multiple comparisons of Tukey’s test, P < 0.05). Error bars represent standard error.

To determine the larvicidal activity of osthole against *T. urticae* and *M. persicae*, we used the same leaf-dipping method to treat the first instar nymphs, and recorded their mortality after 3 days. Both insects showed a dose-dependent mortality trend, with the highest recorded mortality at a concentration of 10 mg/mL osthole (Fig. 3B). The LC_20_ and LC_50_ values for *T. urticae* nymphs were 29% and 8% lower, respectively, than those of adult mites, as shown in Table 3. Similarly, the LC_20_ and LC_50_ values for *M. persicae* nymphs decreased by 85% and 29%, respectively, compared to adults (Table 3). These results suggest that adults are more tolerant to osthole than nymphs.

**Table 3.**
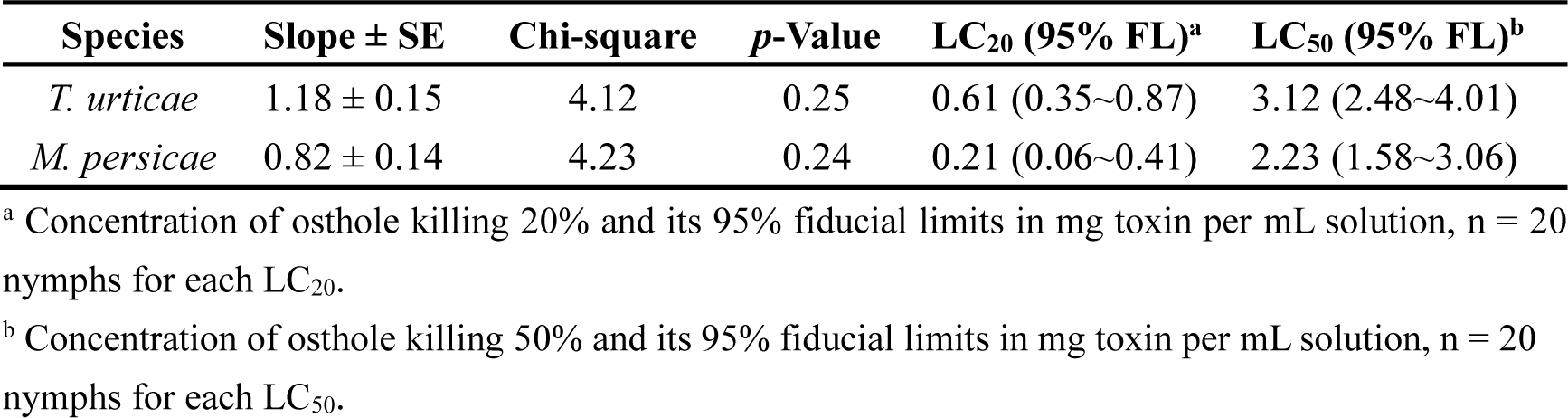
Efficacy of osthole against nymphs of *T. urticae* and *M. persicae*.

| Species | Slope ± SE | Chi-square | *p*-Value | LC<sub>20</sub> (95% FL)<sup>a</sup> | LC<sub>50</sub> (95% FL)<sup>b</sup> |
| --- | --- | --- | --- | --- | --- |
| *T. urticae* | 1.18 ± 0.15 | 4.12 | 0.25 | 0.61 (0.35~0.87) | 3.12 (2.48~4.01) |
| *M. persicae* | 0.82 ± 0.14 | 4.23 | 0.24 | 0.21 (0.06~0.41) | 2.23 (1.58~3.06) |

<sup>a</sup> Concentration of osthole killing 20% and its 95% fiducial limits in mg toxin per mL solution, n = 20 nymphs for each LC<sub>20</sub>.
<sup>b</sup> Concentration of osthole killing 50% and its 95% fiducial limits in mg toxin per mL solution, n = 20 nymphs for each LC<sub>50</sub>.

We tested the toxicity of osthole to *B. dorsalis* adults by feeding them artificial diets with varying concentrations of osthole for 7 days. However, we found no significant difference in survival rates between the experimental and control groups after the feeding period (Fig. 3A). It is worth noting that adult female *B. dorsalis* deposit their eggs within the flesh of soft fruits, and the larvae subsequently hatch and feed inside the fruits (Capinera, 2001). Therefore, it was important to evaluate the toxicity of osthole to *B. dorsalis* adults, but the results suggest that it may not be effective for controlling this pest.

### 3.4 The development and reproduction of *T. urticae* and *M. persicae* were impaired upon osthole exposure

We found that exposure to the LC_50_ concentration of osthole significantly prolonged the larval developmental period of *T. urticae* by 8.32%. Meanwhile, the developmental time of *M. persicae* was significantly increased by 8.92% and 15.9% after treating them with the LC_20_ and LC_50_ concentrations, respectively, when compared to the control group (Fig. 4A). However, the body lengths of both insects remained unchanged (Fig. 4B). The fecundity of adults also significantly decreased after treatment with the LC_20_ and LC_50_ concentrations of osthole, by around 15% for *T. urticae* and 28% for *M. persicae* (Fig. 4C). Exposure to osthole did not cause any adverse effects on *B. dorsalis*, as indicated by the results in Fig. 4C. These results indicate that osthole significantly impairs the development and reproduction of both *T. urticae* and *M. persicae*.

**Figure 4.**
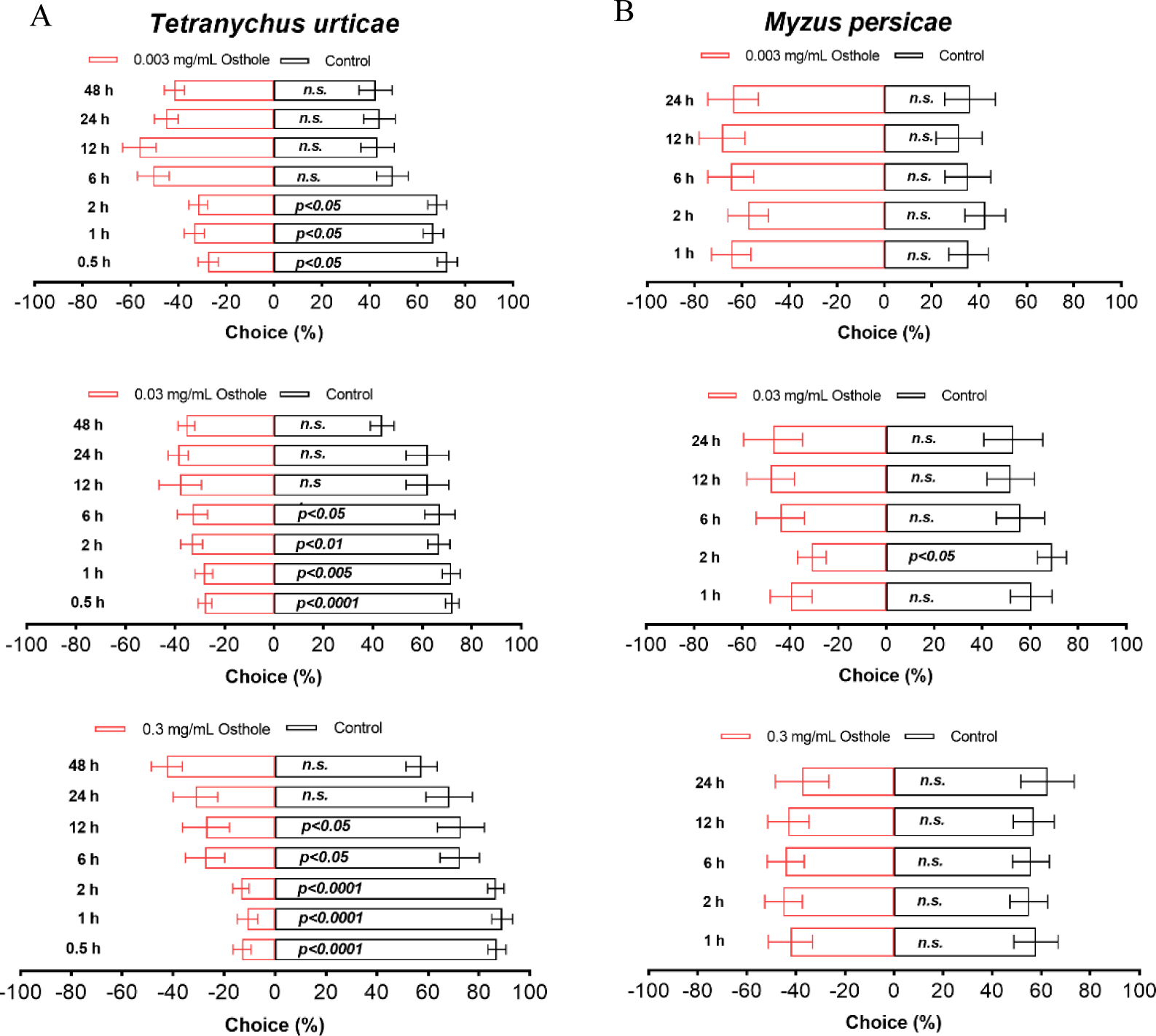
Antifeedant activity of osthole on female adults of *T. urticae* (A) and *M. persicae* (B). The paired sample T-test was used for statistical analysis. All experiments were reproduced 8 times. n. s., not significant.

### 3.5 Osthole present remarkable antifeedant activity against *T. urticae*

We conducted leaf choice bioassays to test osthole for antifeedant activities. Osthole exhibited potent and concentration-dependent antifeedant activities against *T. urticae* (Fig. 5). At 0.5 hours after *T. urticae* release, the number of mites occupying leaves immersed in 0.003 mg/mL osthole was significantly lower compared to control leaves, with this antifeedant effect persisting for a total of 2 hours (∼70% versus ∼30%) (Fig. 5A). Increasing the concentration of osthole to 0.03 mg/mL extended the antifeedant effect for up to 6 hours (Fig. 5A). Similarly, a higher concentration of osthole at 0.3 mg/mL resulted in an even more pronounced and longer response, with the effect lasting for up to 12 hours (Fig. 5A). Notably, in the latter case, the antifeedant effects were even stronger, as a majority of the mites chose control leaves over treated ones (∼80% versus ∼20%) (Fig. 5A). However, no antifeedant activity of osthole was observed for *M. persicae* (Fig. 5B).

**Figure 5.**
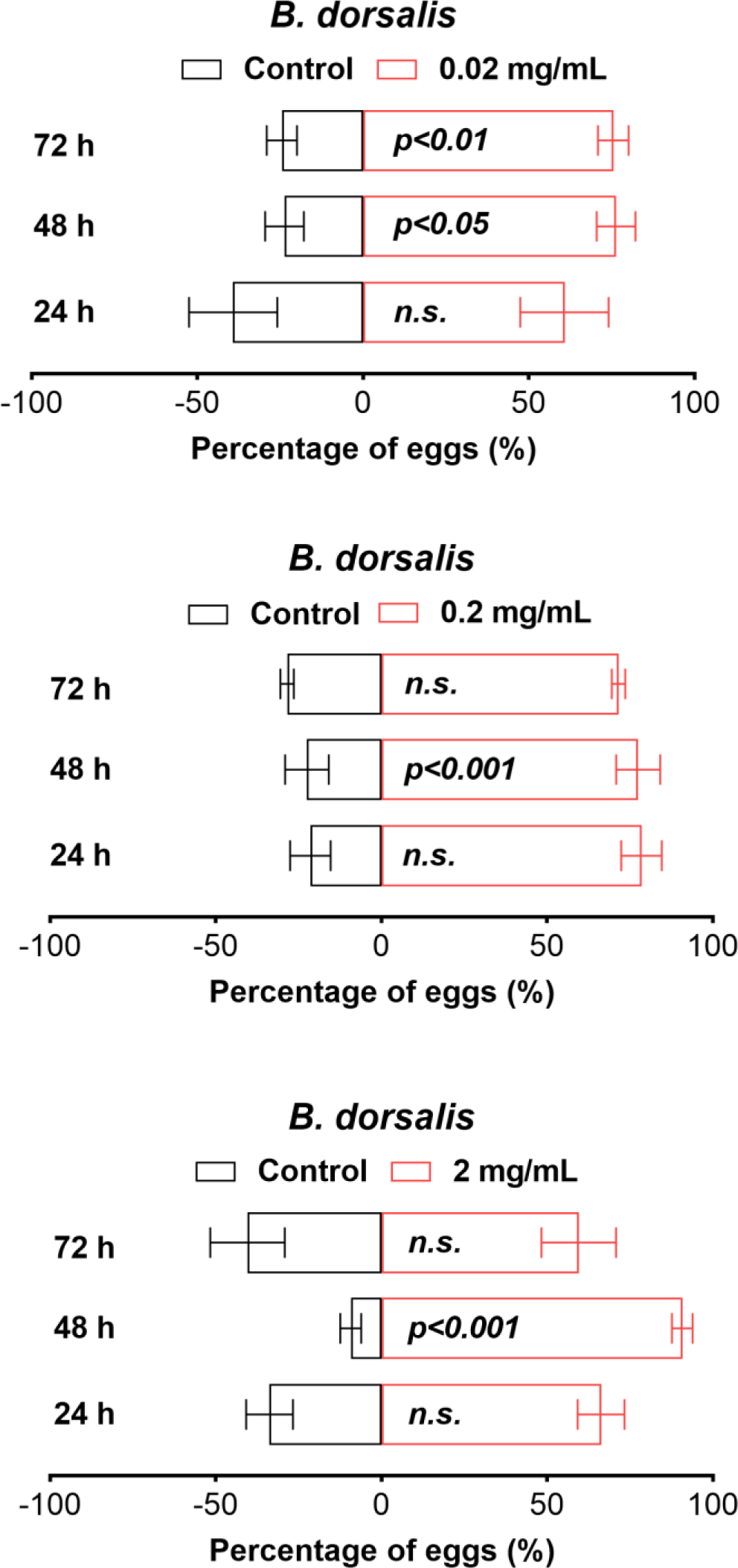
Oviposition attraction of osthole to *B. dorsalis*. The paired sample T-test was used for statistical analysis. All experiments were replicated five times. n. s., not significant.

3.6 Osthole elicited oviposition attraction of *B. dorsalis*

To investigate the oviposition preference of *B. dorsalis* towards osthole, we conducted 2-choice assays within a transparent cage. Our results indicated that female flies displayed a tendency to lay their eggs in mango juice containing 0.02 mg/mL of osthole after 48 hours, and this preference was still observable at the 96-hour mark (Fig. 6). Remarkably, the percentage of eggs laid in the presence of osthole (at the 0.02 mg/mL concentration) was around 75%, compared to a mere 25% in its absence (Fig. 6). However, when we increased the concentration of osthole to 0.2 and 2 mg/mL, this preference disappeared by the 96^th^ hour (Fig. 6).

## 4 Discussion

Previous researches have reported that CMC contains 67 types of coumarins, though the majority of their individual contents remains unknown (Sun et al., 2020). In our study, we quantified 11 types of coumarins, including 3 simple coumarins, 6 linear furanocoumarins, and 2 angular furanocoumarins. Osthole was found to be the most abundant type of coumarin present (11.04 mg/g), which is consistent with previous findings (Table S1) (Chen et al., 2009; Gao et al., 2013). Imperatorin content in CMC is reported to range from 2-9 mg/g, while the contents of isopimpinellin and 5-methoxypsoralen range from 1-4 mg/g (Table S1) (Chen et al., 2009; Gao et al., 2013; Wang et al., 2010). In our study, we found the contents of imperatorin, isopimpinellin, and 5-methoxypsoralen in CMC from Wudi (Shandong province) to be 3.41, 0.64, and 0.45 mg/g, respectively. As *C. monnieri* is widely cultivated in various regions of China, our findings suggest that the contents of coumarins in CMC exhibit quantitative variation depending on the geographic region. Furthermore, our study is the first to report the contents of the remaining 7 coumarins in CMC.

In CMC, coumarins are considered the primary bioactive compounds based on previous research studies (Cai et al., 2000; Liu et al., 1999; Sun et al., 2021; Wu et al., 2017). In our study, we examined the toxicity of osthole and four other coumarins which are found in CMC against *T. urticae*. Interestingly, we discovered that all of the coumarins exhibited similar toxicity towards the mites. Moreover, scoparone, which is also a well-known coumarin with potent acaricidal activity against mites (Zhou et al., 2022), demonstrated similar toxicity towards *T. urticae* as well. These findings suggest that osthole, believed to be the major bioactive compound in CMC, is likely responsible for the observed effects primarily due to its high content in CMC.

Although osthole is known for its beneficial biological and pharmacological activities, there is limited information regarding its insecticidal properties (Sun et al., 2021). Previous studies have identified its toxicities against oriental armyworm (*Mythimna separate*), diamondback moth (*Plutella xylostella*), green peach aphid (*M. persicae*), two-spotted spider mite (*T. urticae*), and mosquitoes (*Culex pipiens pallens* and *Aedes aegypti*) (Li et al., 2021; Ren et al., 2020; Ren et al., 2021; Wang et al., 2012; Yan et al., 2021). In our study, the LC_50_ value against *T. urticae* decreased from 4.39 to 2.49 mg/mL over a period of 5 days after treatment with osthole, while the LC_50_ value against *M. persicae* decreased from 4.56 to 2.95 mg/mL. These results are more than 2-fold higher than the LC_50_ value of osthole against *T. cinnabarinus* (the red form of *T. urticae*) reported previously, indicating potential differences in tolerance towards osthole between the two forms (Li et al., 2021; Ren et al., 2020). Additionally, the LC_50_ values of osthole against third-instar larvae of *A. aegypti* and *C. p. pallens* were determined as 0.013 mg/mL (Wang et al., 2012; Yan et al., 2021), suggesting that osthole may exhibit more potent insecticidal activity against mosquitoes.

Previous studies have indicated that coumarins have a detrimental effect on the growth, development, and reproduction of insects (Berenbaum, 1983; Pavela and Vrchotová, 2013). To evaluate the toxicity of osthole towards *M. persicae*, *T. urticae*, and *B. dorsalis*, we conducted exposure experiments using various concentrations of osthole based on LC_20_ and LC_50_ values. Our findings demonstrated that osthole treatment prolonged the developmental time of *M. persicae* and *T. urticae*, while also impairing their fecundity. These results support previous reports indicating that osthole can prolong the developmental time of *M. separate* (Li et al., 2021).

Prior research has suggested that coumarins act as potent antifeedants and oviposition deterrents against insects (Poudel et al., 2015; Poudel and Lee, 2016; Stevenson et al., 2003; Tabashnik, 1987). However, it is unknown whether osthole can function as an antifeedant and oviposition deterrent. Therefore, our study is the first to report on the antifeedant effects of osthole against *T. urticae* and its unexpected oviposition attraction property towards *B. dorsalis*. To the best of our knowledge, no published report has described coumarins with oviposition attraction activity. *B. dorsalis* is a highly polyphagous fruit fly species that infests around 450 plant species worldwide, including *Citrus* spp., which contain osthole (Clarke et al., 2005; Zhang et al., 2002). Thus, the oviposition attraction activity of osthole towards *B. dorsalis* and its lack of toxicity might be attributed to the insect’s strong detoxification capacity, developed through a long co-evolutionary process with *Citrus* plants. The preference of *B. dorsalis* for osthole makes it a suitable attractant for use in the control of *B. dorsalis*.

The cost of botanical pesticides has been a barrier to their commercial success (Isman, 2006; Isman, 2020). Excitingly, our recent research has demonstrated the high effectiveness of *C. monnieri* flower strips in supporting natural enemies, suppressing aphid populations, and reducing insecticide use across wheat-maize rotations, cotton crops, and apple orchards (Yang et al., 2020; Yang et al., 2022; Zhang et al., 2022). The Chinese government has already authorized the use of *C. monnieri* flower strips in pest management, providing additional optimism for a reduction in osthole usage costs.

## 5 Conclusion

In conclusion, our research has elucidated that osthole is the predominant coumarin present in CMC and displays efficient insecticidal activity against *T. urticae* and *M. persicae*. We have provided novel evidence demonstrating that osthole acts as an effective feeding deterrent for female *T. urticae* adults, and conversely, as a strong attractant for female *B. dorsalis* adults. Consequently, our findings suggest that osthole has significant potential as a botanical insecticide for the management of *M. persicae*, *T. urticae* and *B. dorsalis*.

## Acknowledgements

This work is supported by grants from Agricultural Science and Technology Innovation Project of Shandong Academy of Agricultural Sciences [grant number CXGC2023F04 and CXGC2023A24] and the Taishan Scholars Program.

---

**Table S1.**
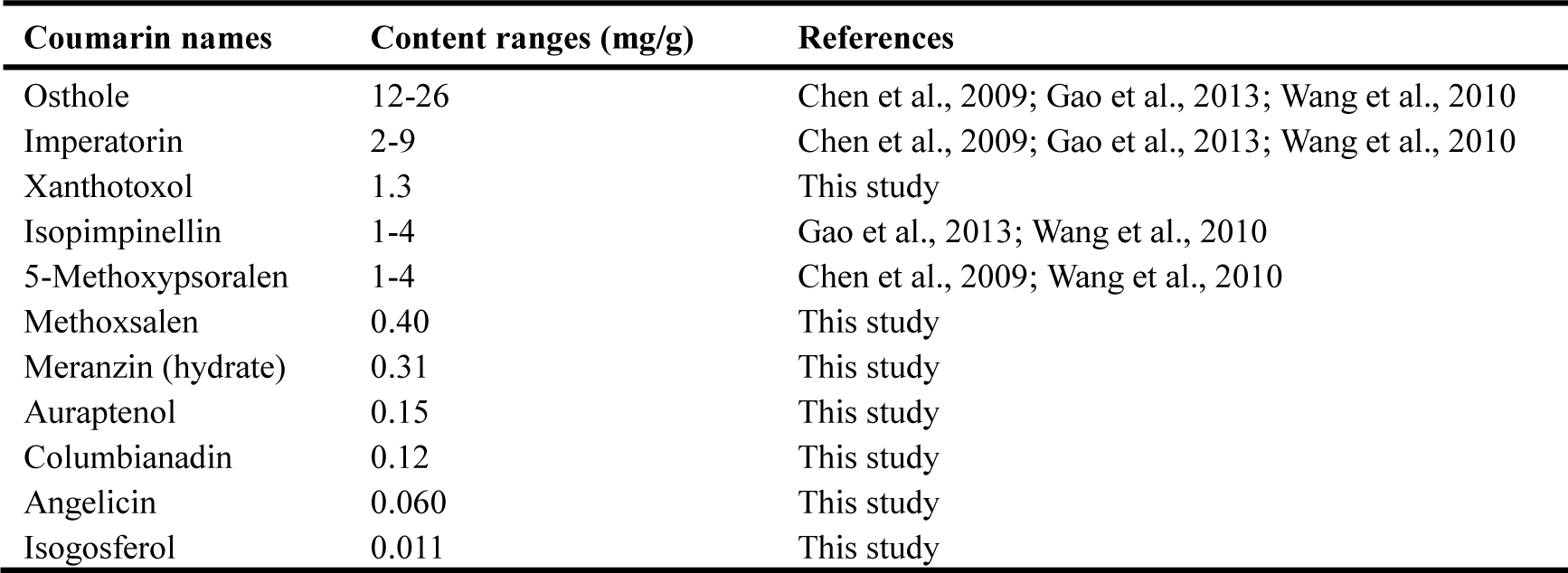
Contents of coumarins in CMC.

