## Supplementary material for "Contact toxicity, antifeedant activity and oviposition preference of osthole against agricultural pests": Table S1

Table S1 Contents of coumarins in CMC.

| **Coumarin names** | **Content ranges (mg/g)** | **References** |
| --- | --- | --- |
| Osthole | 12-26 | Chen et al., 2009; Gao et al., 2013; Wang et al., 2010 |
| Imperatorin | 2-9 | Chen et al., 2009; Gao et al., 2013; Wang et al., 2010 |
| Xanthotoxol | 1.3 | This study |
| Isopimpinellin | 1-4 | Gao et al., 2013; Wang et al., 2010 |
| 5-Methoxypsoralen | 1-4 | Chen et al., 2009; Wang et al., 2010 |
| Methoxsalen | 0.40 | This study |
| Meranzin (hydrate) | 0.31 | This study |
| Auraptenol | 0.15 | This study |
| Columbianadin | 0.12 | This study |
| Angelicin | 0.060 | This study |
| Isogosferol | 0.011 | This study |
